# Metabolic changes that allow *Plasmodium falciparum* artemisinin-resistant parasites to tolerate the oxidative stress

**DOI:** 10.1101/2023.10.21.560494

**Authors:** Alejandro David Bonive-Boscan, Héctor Acosta, Ascanio Rojas

## Abstract

1.

**Background:** Artemisinin-based treatments (ACTs) are the first therapy currently used to treat malaria produced by *Plasmodium falciparum*. However, in recent years increasing evidence shows that some strains of *P. falciparum* are less susceptible to ACT in the Southeast Asian region.

**Materials & Methods:** A data reanalysis of several omics approaches currently available about parasites of *P. falciparum* that have some degree of resistance to ACT was carried out. The data used was from transcriptomics and metabolomics studies. One mitochondrial carrier of the parasite possibly involved in the mechanisms of tolerance to oxidative stress was modelled and subjected to molecular dockings with citrate and oxoglutarate.

**Results:** An increase in glutathione production was detected, changing the direction of the flux of metabolites in the tricarboxylic acid cycle, and boosting glucose consumed. The models of the mitochondrial carrier, called PfCOCP, show that it may be important in transporting citrate and oxoglutarate from the mitochondrial matrix to cytosol. If so, it may allow the parasite to tolerate the oxidative stress produced by artemisinin.

**Conclusions:** This *in-silico* analysis shows that *P. falciparum* may tolerate the artemisinin’s oxidative stress through metabolic changes not reported before, showing the need for further research on the many metabolic aspects linked to this phenotype.

## 2 Background

Malaria causes an estimated 229 million cases and 409.000 deaths in 2019; the majority of these were produced by *Plasmodium falciparum* [1]. From 2001, the treatments based on Artemisinin (ACTs) have been primary drugs used against malaria produced by *P. falciparum*. Artemisinin (ART) affects the parasite in different ways: interacting with and damaging proteins, inhibiting the proteasome, and producing *Reactive Oxygen Species* (ROS)(artemisinin resistance have been reviewed elsewhere [2]). Those effects produce the death of parasites interfering with several metabolic pathways causing systemic damage.

Despite the multilevel damage produced by the drug in 2008, some evidence suggested that the ACTs were losing their efficacy against *P. falciparum* in the Greater Mekong Subregion (GMS) in Southeast Asia [3]. Further studies show that mutations in the protein 13 of *P. falciparum* are the main molecular marker of resistant parasites [4], and many mutations have been identified worldwide, several associated with an increase in the number of surviving parasites in blood [5]. However, the resistant parasites show other changes besides mutations in Pfkelch 13. Some parasites obtained *in vitro* show an increase in tolerance to artemisinin lacking mutations in PfKelch 13 [6]. One of the metabolic changes shown by Dd2 resistant strains is an increase in the oxidative stress response [7]. Also, resistant strain Cam3.II with C580Y or R539T mutations of PfKelch13 show higher concentrations of Glutathione (GSH) and γ-glutamylcysteine, both metabolites are directly associated with oxidative stress response [8]. Nowadays a lot of evidence show artemisinin resistance is a very complex process that may be very different depending genetic background [2].

A pathway that shows high plasticity in *P. falciparum* is the Tricarboxylic acid (TCA) cycle. This pathway is used by the parasite for different roles during its life cycle and is dispensable for ATP production when the parasite is in the intraerythrocytic development cycle (IDC) [9]; an asexual cycle composed by the ring, trophozoite and schizont stages. Furthermore, in artemisinin-resistant parasites, the TCA cycle is very important even though it is not vital for ATP production during IDC. The resistant strains have shown more plasticity in the TCA cycle based on metabolic network reconstructions [10].

An aspect that has been less studied is the mitochondrial carriers that transport metabolites involved in the TCA cycle between mitochondrial matrix and cytosol. Among those, the *P. falciparum* mitochondrial dicarboxylate-tricarboxylate carrier (PfDTC) is the most studied [11]. Likewise, a general approach has been used to study the transport activities of 12 *P. falciparum* mitochondrial carriers [12]. Mitochondrial carriers may be very important to allow *P. falciparum* to change in metabolites fluxes through matrix membrane, allowing TCA cycle changes in different parasite stages. In the yeast *Saccharomyces cerevisiae*, one mitochondrial carrier (YHM2) has been involved in the oxidative stress response [13]. In this study a probable ortholog of the yeast’s mitochondrial carrier in *P. falciparum* was identified (PlasmoDB ID PF3D7_1223800). This protein may play a similar role in *P. falciparum* contributing to the oxidative stress response.

The present research will re-analyze part of the omics data about *P. falciparum* artemisinin-resistant parasites. The study proposes a metabolic model that explains possible mechanisms that allow the parasite to tolerate the oxidative stress produced by the artemisinin, one of the drug’s effects that has been previously reported [14]. The study will focus on three essential routes for the parasite: Glycolysis, TCA cycle, and glutathione production. Additionally, the mitochondrial carrier PF3D7_1223800 will be studied *in silico*, based on the recognized importance of this carrier in the oxidative stress response induced by the drug. These approaches can show new possible therapeutic targets affecting the oxidative stress response in *P. falciparum*.

## 3 Methods

### 3.1 Metabolic model of mechanisms to tolerate the oxidative stress produced by the artemisinin

We use the transcriptomic profiles [15] on more than 1000 clinical samples of *P. falciparum* in GMS and other places. These data contain 999 protein transcripts positively and negatively statistically correlated with clinical samples that show an increase in the tolerance to artemisinin. We posited that differences in the transcripts’ abundance imply differences in the protein quantity. Furthermore, the metabolomic, proteomic, and transcriptomic changes using the strain Cam3.II carrying C580Y, and R539T mutations in PfKelch 13 were taken into account [8, 16]. Those data obtained by different sources were used to construct a theoretical metabolic model of resistant strains; the metabolic pathways of *P. falciparum* were obtained from The *Malaria Parasite Metabolic Pathways* website (MPMP, [17]).

### 3.2 Modelling a mitochondrial carrier possible involved in the oxidative stress response in *P. falciparum*

The putative citrate/oxoglutarate transporter (PF3D7_1223800) was renamed as Citrate/Oxoglutarate Carrier Protein of *P. falciparum* (PfCOCP). This protein was modelled *in silico* using Phyre2 [18] and SwissModel [19] to obtain the carrier in the two different conformations that have been previously described [20]. Phyre2 automatically uses different templates that have a good fit with the sequence supplied. SwissModel follows the same process, but the model obtained was less accurate than the one obtained with Phyre2. Finally, we used Phyre2 to obtain the carrier in Cytosolic (C) state. Phyre2 does not allow the use of a specific template. Consequently, we used SwissModel to explicitly obtain the carrier in Matrix (M) state, using the template obtained [21](PDB ID: 6GCI,) of the ADP/ATP carrier from the thermotolerant fungus *Thermothelomyces thermophila* inhibited by Bongkrekic acid.

The models obtained were refined using GalaxyWeb [22]; twice for the model in C state and once for the model in M state. The models were refined until no improvements were found in the Rama favoured parameter, which shows how favoured the protein is, based on Ramachandran’s plot [23].

Molecular dockings were carried out with the refined models. This was done using two substrates obtained from the ZINC database [24]: citrate (ZINC895081) and oxoglutarate (ZINC1532519). The dockings were made using SwissDock [25].

UCSF Chimera 1.13.1[26] was used to visualize protein models and dockings obtained from the different servers. In addition, using the Chimera function, *FindHBond* helped us identify which residues possibly interact with citrate and oxoglutarate in the protein.

## 4 Results

### 4.1 Metabolic model of mechanisms to tolerate the oxidative stress produced by the artemisinin

We mainly used the transcriptomic and metabolomic data obtained from *P. falciparum* artemisinin-resistant strains [8, 15, 16] to propose a metabolic model that shows an increase in the glycolytic path based on an increase in transcript levels of hexokinase (HK, PF3D7_0624000), phosphoglucose isomerase (PGI, PF3D7_1436000), phosphofructokinase (PFK, PF3D7_0915400), and phodphoglicerate kinase (PGK, PF3D7_0922500) in clinical samples [15].These are highlighted in **Figure 1**.

**Figure 1:**
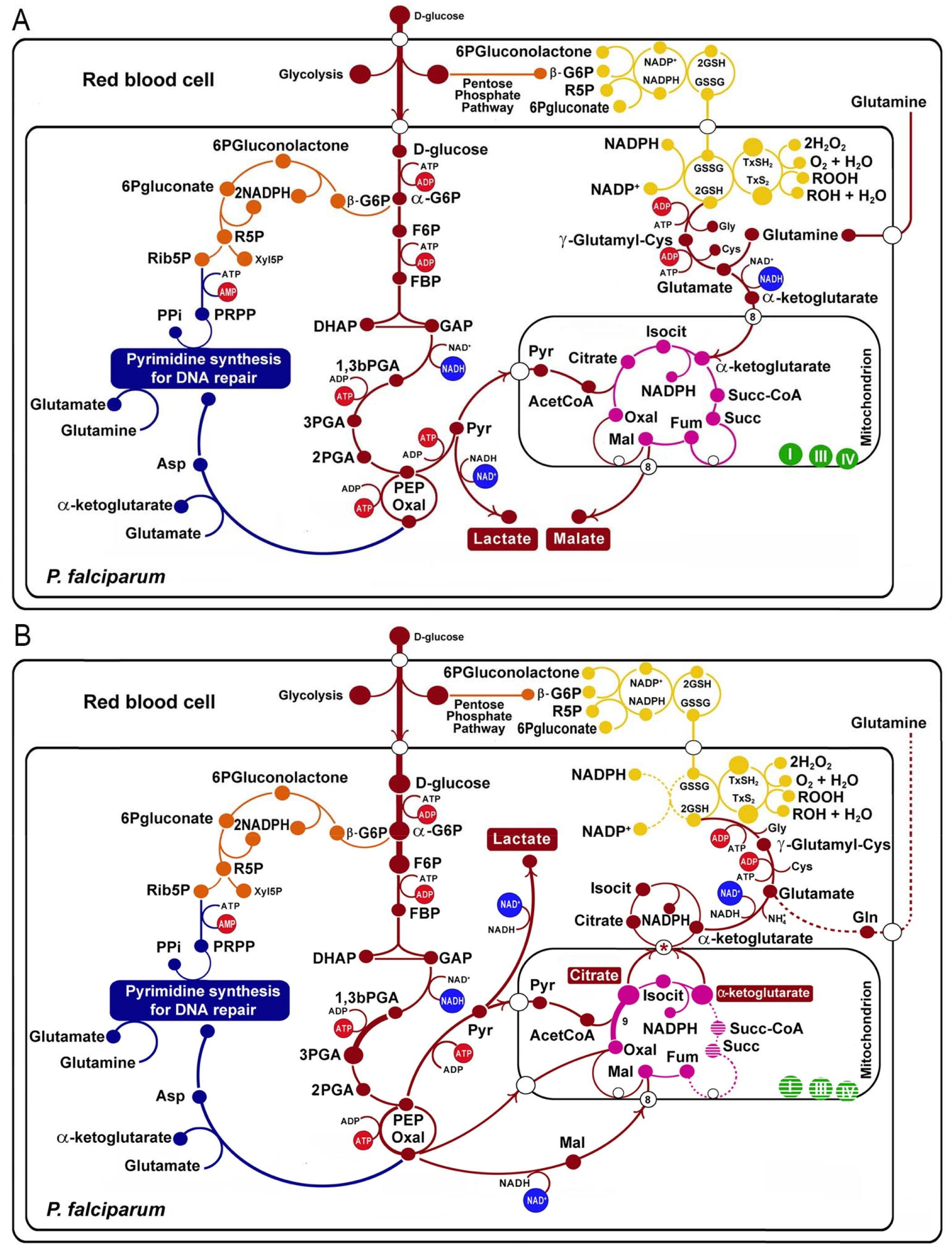
Metabolic changes in artemisinin resistant strains of *P. falciparum* (**B)** compared with non-resistant strains (**A**), the wide light red arrows indicate the principal flux of metabolites across the Glycolysis, TCA cycle and Glutathione production. The proteins transcripts positively correlated with the resistance are indicate with a thick line, negatively correlated are indicate with a dotted line, according with [15]. Glycolytic pathway in red: -G6P: alpha glucose-6 phosphate; F6P: fructose-6 phosphate; FBP: fructose 1,6-biphosphate; DHAP: dihydroxyketonephosphate; GAP: glyceraldehyde 3-phosphate; 1,3bPGA: glycerate 1,3 biphosphate; 3PGA: 3-phosphoglycerate; 2PGA: 2-phosphoglycerate; PEP: phospho*enol*pyruvate. Tricarboxylic acid cycle in violet: OXA: oxalacetate; Isocit: isocitrate; Succ-CoA: succinic Coenzyme A; Succ: succinate.Pentose phosphatepathway in orange: -G6P: beta glucose-6 phosphate; R5P: ribulose-5 phosphate.Purine and pyrimidine synthesis in blue: Xyl5P: xilulose-5 phosphate; Ribo5P: ribose-5 phosphate; DHA: dihydroxycetone; PRPP: phosphoribosyl pyrophosphate. Oxidative stress branch in yellow: GSH2: glutathionereduced; G2S2; GXS2; GXSH2. The asterisks indicate the PfCOCP (a citrate/oxoglutarate mitochondrial carrier).

The tricarboxylic acid (TCA) cycle shows marked differences in the resistant strains, also based on transcripts levels changes. For example, citrate synthase (PF3D7_1022500, PF3D7_0609200) is upregulated and oxoglutarate dehydrogenase (PF3D7_1320800), succinate dehydrogenase (PF3D7_1010300), and succinate CoA ligase (PF3D7_1108500) are downregulated. These changes in the transcript levels may be related to changes in the metabolites influx and efflux from the mitochondrion, specifically, this can promote the efflux of oxoglutarate and influx of malate, putatively; the transport can occur using the malate/oxoglutarate mitochondrial carrier of *P. falciparum* (PfDTC, PF3D7_0823900). For its part, citrate is overproduced and this can imply its acumulation in the mitochondria matrix. This was determined for non-resistant parasites in metabolomic analysis, besides, resistant parasites show less citrate acumulation in drug presense [16], a fact that may imply that resistant parasites are better transporting citrate to cytosol. Citrate can be transported to cytosol using the carrier PF3D7_1223800; this protein is a homolog of the citrate/oxoglutarate mitochondrial carrier of *Saccharomyces cerevisiae* [13].

On the other hand, metabolomic analyses were made in *P. falciparum* Cam3.II strains with R539T and C580Y mutations in PfKelch 13 (Cam3.II^R539T^ and Cam3.II^C580Y^) show that Glutathione and γ-glutamylcysteine are more abundant in these resistant strains [8]. This evidence was used to propose that glutathione production is overexpressed in some resistant strains (**Figure 1**). Moreover, the citrate and oxoglutarate transported to the cytosol (mentioned above) can contribute to glutathione production.

For the reduction of glutathione and thioredoxin, the parasite needs NADPH. **Figure 1** shows that diverse NADP(H) oxidoreductases carry on the reaction. Isocitrate dehydrogenase may be the primary source of NADPH in the cytosol. In fact, increased utilization of glucose by the glycolytic pathway and not by Pentoses Phosphate pathway (PPP), together with a decrease in the levels of transketolase transcripts (PF3D7_0610800) of PPP [15], suggest that PPP is less relevant for NADPH production in the resistant strains.

Thus far, all the changes proposed agree with an increase of the oxidative stress tolerance in some resistant strains in compared with sensible ones.

### 4.2 Modelling a mitochondrial carrier possibly involved in the oxidative stress response in *P. falciparum*

The protein PfCOCP was modelled and subjected to molecular dockings with citrate and oxoglutarate. PfCOCP shows 6 transmembrane regions (**Figure 2 A**) formed by α-helixes. These regions are going to be named H1, H2, H3, H4, H5, and H6 following other research done on these carriers (reviewed in [20]). There are also 3 domains in the carrier’s mitochondrial matrix side that connect the different transmembrane regions, identified as H12, H34, and H56, which are also found in the works previously cited. These carriers have two different conformations: opened to Cytosol (C state) and opened to Matrix (M state). PfCOCP was modeled in two conformational forms, cytosol(C) and matrix (M) state (**Figure 3**), using Phyre2 and SwissModel, respectively.

**Figure 2:**
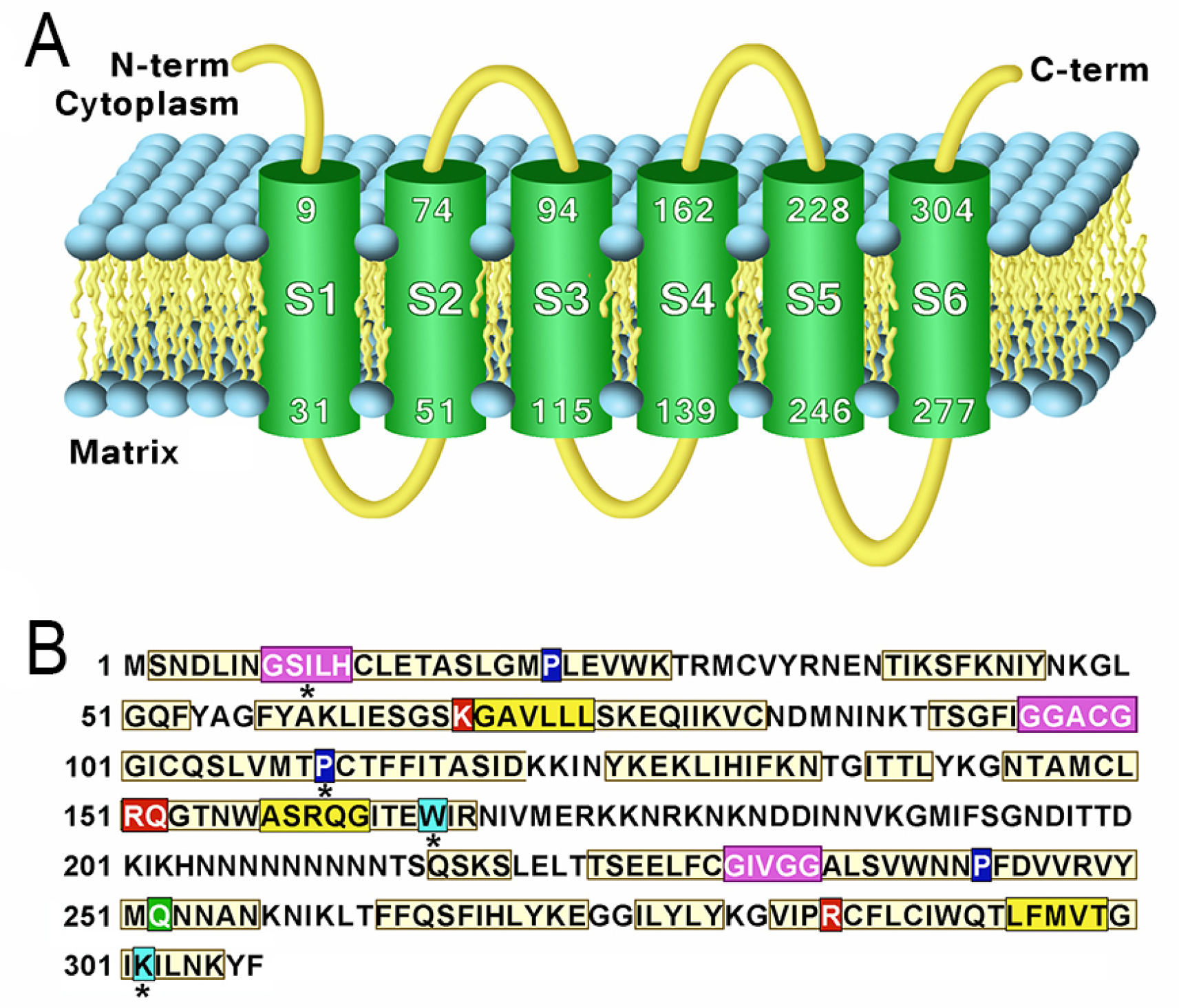
A transmembrane regions of the model obtained using Phyre 2 of *P. falciparum* mitochondrial Citrate/Oxoglutarate carrier (PfCOCP) in C state. **B** Sequence features found in (PfCOCP): GxxxG motif (violet), πxxxπ(yellow), Pxx[DE]xx[RK] (proline in blue), Q braces (green), Y braces (cyan) and substrate binding sites (red). Asterisks (*) show motif that do not match perfectly with motif reported in literature.

**Figure 3:**
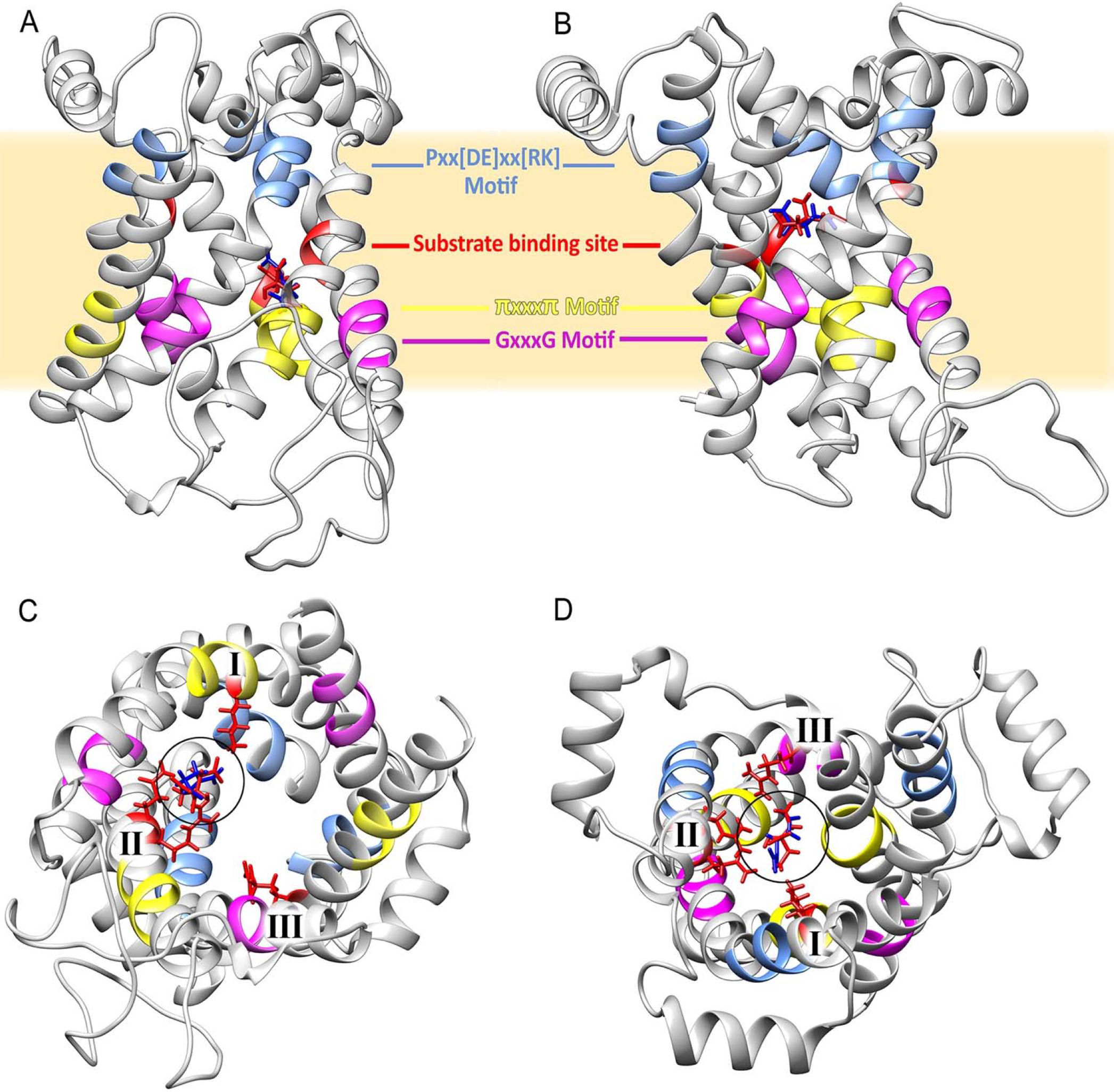
Structure of Citrate/Oxoglutarate mitochondrial carrier of *P. falciparum* modelled in Cytosol (**A**) and Matrix (**B**) state. Some of the sequence features are highlighted in both structures. (**C**) show the protein in C state viewed from cytosol (**D**) showed the protein in M state viewed from matrix, chain of residues that interact with substrates are showed in red and the number of biding sites (I, II and III) are showed in both cases.

We found several motifs normally identified in mitochondrial carriers (**Figure 2 B**). GxxxG was found in odd segments, namely in H3 (**G**GAC**G**) and H5 (**G**IVG**G**), and tentatively in H1 (**G**SIL**H**). The latter agrees with the location if we highlight the H3’s and H5’s motif in the models (**Figure 3 A** and **B**). This motif and the other regions that do not totally agree with the motif postulated in the existing literature were noted with an asterisk (*) in **Figure 2 B**. πxxxπ were found in even segments: in H2 (**G**SKG**V)**, H4 (**T**MWA**S)**, and H6 (**T**LFM**V**). Motif [PS]x[DE]xx[RK] were found in odd segments: H1 (**P**L**E**VW**K**), H5(**P**F**D**VV**R**), and tentatively in H3(**P**CTFFI). Proline residues found in [PS]x[DE]xx[RK] are important for the torsion of the transmembrane domains and they allow mitochondrial carriers to change between both states (C and M) [20]. One Q brace was identified in H5 (**P**F**D**VV**R**VYM**Q**) and two tentative Y braces in (**Figure 2 B**). These motifs have been previously described in other mitochondrial carriers [20]. On the other hand, the [FY][DE]x[RK] motifs could not be identified, which shows that these motifs are less conserved in mitochondrial carriers [20].

Molecular dockings with citrate (red) and oxoglutarate (blue) are shown in **Figure 3**. Gibbs free energy (ΔG) for citrate and oxoglutarate in C state are -18.37 and -24.85 kcal/mol; for M state, they are -21.33 and -31.07 kcal/mol, respectively. This suggests that PfCOCP the M state better interacts with both substrates. PfCOCP has three putative binding sites that interact with substrates; these are usually named I, II, III and are located in H2, H4, and H6 transmembrane domains of mitochondrial carriers [27]. The residues that possibly interact with substrates were identified in C and M states. This process allows us to propose the three sites for substrate interaction, site I in H2 (Lys 67), site II in H4 (Arg 151, Gln 152), and site III in H6 (Arg 262). The fact that site I has a Lys, site II is composed of Arg-Gln, and site III has an Arg, agrees with the substrate-binding sites of other mitochondrial carriers: namely, succinate carrier (PDB ID: FC1P) and citrate/malate carrier (TP1P), both of *S. cerevisiae* [27].

Only in the M state of PfCOCP, the three substrate binding sites interact with citrate or oxoglutarate; besides that, in the C state, only sites I and II interact with substrates (**Figure 3 C** and **D**). These differences also suggest that PfCOCP is better at interacting with substrates in the M state than the C state. The differences in ΔG, substrate binding sites and the accumulation of matrix citrate and oxoglutarate allow us to propose that PfCOCP transport citrate and oxoglutarate from matrix to cytosol as is showed in **Figure 1**.

## 5 Discussion

The resistance of *P. falciparum* to artemisinin has been studied extensively. This resistance has been attributed to different mechanisms. Different mechanisms used by *P. falciparum* to tolerate the artemisinin effects may be present in the same parasites, actually, the data used in this study are from PfKelch 13 mutant parasites, a fact that has been associated with a reduction in hemoglobin intake and heme production [2].

This study’s metabolic pattern is consistent with an increase in oxidative stress response, which has been previously reported [7, 14]. Our model focused on three metabolic routes: Glycolysis, TCA cycle, and Glutathione production.

The first observation is the increase in the expression of the first enzymes of glycolysis: hexokinase (HK), phosphoglucose isomerase (PGI), and phosphofructokinase (PFK), as well as phosphoglycerate kinase (PGK). Interestingly, two of these enzymes (HK and PFK) are described as enzymes with a high flux coefficient in the glycolytic kinetic model of *P. falciparum* [28]. We believe that the overexpression of these enzymes could be reflected in an increase in glucose consumption by the parasite. Additionally, overexpression of PGK could favor a higher percentage of glucose processed by the glycolytic pathway versus the pentose phosphate pathway (PPP). Moreover, expression levels of both Glucose-6 phosphate dehydrogenase and 6-phosphogluconate dehydrogenase remain similar to those non-resistant parasites. Likewise, in these data transketolase is under expressed [15]. However, the route should be operative since the parasite needs the production of Ribulose-5P to synthesize PRPP, which is essential for the synthesis of pyrimidines.

The use of this glucose by the glycolytic pathway will lead to phosphoenolpyruvate (PEP) formation, which is at a metabolic bifurcation point. One part of glucose is transformed into pyruvate by a pyruvatekinase (PK), providing the first net gain of ATP, while the other part of PEP can be transformed into oxaloacetate by a phosphoenolpyruvate carboxylase (PEPC) or phosphoenolpyruvate carboxykinase (PEPCK), which provides the second net ATP gain. These three enzymes (PK, PEPC and PEPCK) do not show expression changes in transcriptomic data obtained from clinical samples [15], but in a recent research with artemisinin resistant strains Dd2 and Cam3.II of *P falciparum*, high levels of PEPCK transcripts have been found [16].

Another interesting change occurs in the TCA cycle where an increase in the expression of citrate synthase together with the under expression of oxoglutarate dehydrogenase, succinate-CoA ligase and succinate dehydrogenase, would point to the functionality of only the first steps of this cycle. One consequence of this is the accumulation of citrate and/or oxoglutarate on the mitochondria and for this, both acetyl-CoA and oxaloacetate must be supplied continuously. Thus, a metabolomic study on the artemisinin-resistant strain Cam3.II^C580Y^ showed high concentrations of citrate, which would support the previously described model [16]. We gather that the function of the TCA cycle in these resistant strains is similar to the function observed in *P. falciparum* parasites with knockout gene of oxoglutarate dehydrogenase, which would force malate to enter the mitochondria to supply oxaloacetate via MQO and oxoglutarate to exit from the organelle [9].

From the above, it follows that a constant supply of oxaloacetate for citrate synthase would require an efficient MQO. However, the decrease in the expression of components of complexes III and IV of the electron transport chain would negatively affect this supply of oxaloacetate resulting in a possible accumulation of malate. Recent metabolomic evidence suggests that malate can accumulate in Cam3.II^C580Y^ [16]. Another alternative could be the direct entry of the oxaloacetate, thus avoiding the possible low functionality of MQO. In this sense, the use of mitochondrial malate/oxoglutarate carrier of *P. falciparum*, called the dicarboxylate/tricarboxylate carrier (PfDTC) [11], would be an alternative. This transporter proved to be efficient in transporting the oxoglutarate/malate pairs and oxoglutarate/oxaloacetate across liposome membranes, so oxaloacetate could be transported to the mitochondrial matrix and form citrate even if malate accumulates in the organelle.

Different evidence shows that Cam3.II^R539T^, Cam3.II,^C580Y^ and Dd2 strains of *P. falciparum* are obliged to constantly synthesize glutathione during infection to combat the oxidative stress produced by artemisinin [7, 8]. In this sense, the oxoglutarate that accumulates in the mitochondria should leave and contribute to the demand for glutathione by the parasite. We also suggest that the mitochondria’s citrate could have the same fate as oxoglutarate to form glutamate in the cytosol. In this regard, during the investigation, we found that PfCOCP, a mitochondrial citrate/oxoglutarate carrier, has an increase in the level of transcripts of mutant strain PfKelch 13 of *P. falciparum* [15].

We think that this protein may play an important role in the transport of citrate and oxoglutarate from the matrix to the cytosol. In this context, our modeling, together with the studies of molecular couplings with citrate and oxoglutarate, show that PfCOCP could transport citrate more efficiently from the mitochondrial matrix to the cytosol than vice versa (Figure 3), a fact that agrees with the role we assigned to this transporter.

PfCOCP was proposed as a putative ortholog in *P. falciparum* of the YHM2 protein of *S. cerevisiae*, a mitochondrial citrate/oxoglutarate transporter that was associated with the response to oxidative stress in yeast [13]. PfCOCP may have played this role in *P. falciparum*. It should be noted that the citrate transported to the cytosol needs two enzymes to transform into oxoglutarate: aconitase and isocitrate dehydrogenase. *P. falciparum* aconitase has been localized to mitochondria and another subcellular region, possibly the cytosol or the digestive vacuole [29]. On the other hand, isocitrate dehydrogenase, apart from its location in the mitochondria, may have another subcellular location [30], furthermore in *P. knowlesi* most of the isocitrate dehydrogenase activity has been measured in the cytosolic fraction [31]. Further research is needed to clarify whether both enzymes are in the cytosol.

In the cytosol, the oxoglutarate can be transformed in glutamate via three enzymes: two glutamate dehydrogenases (GDH1, GDH3) and the aspartate aminotransferase (AspAT). GDH1 and GDH3 use NADPH or NADH, respectively [32]. Of those two, we think that *P. falciparum* use GDH3 in artemisinin-resistant strains based on three factors: first, to tolerate the oxidative stress the parasite may need NADPH for recycling glutathione and thioredoxin; second, metabolomic data obtained from resistant strains show an increase in NAD^+^ concentration [8], which may be a consequence of GDH3 overuse; and third, in non-resistant parasites GDH3 has a higher expression than GDH1 or AspAT [33]. Notably, recent evidence obtained from proteomic shows that GDH1 and GDH3 are overexpressed in trophozoites of Cam3.II^R539T^ and Cam3.II^C580Y^, as well as high transcripts of GDH1 in Dd2 resistant strain [16].

This glutathione synthesis is essential for the blood stages of *P. falciparum* [34]; besides, Cam3.II^R539T^ and Cam3.II^C580Y^ strains show higher amounts of glutathione and glutamyl-cysteine (glutathione precursor) compared with unmutated strains [8]. Moreover, in a recent proteomic approach, several enzymes involved in glutathione production show high values in those strains [16]. In clinical samples, transcriptomic studies have shown a decrease in the expression of glutathione reductase [15], an enzyme necessary for the reduction of glutathione during oxidative stress. This was previously reported in non-resistant parasites showing that if the parasite produces more glutathione, there are less recycling of it and more efflux of oxidized glutathione to RBC [34]. We must also note that Dd2 *in vitro* artemisinin-resistant strains overexpress GR [7].

This model implies that glucose may be the primary source of carbon for TCA cycle, agreeing with the mutants obtained by Ke H, Lewis IA, Morrisey JM, McLean KJ, Ganesan SM, Painter HJ, Mather MW, Jacobs-Lorena M, Llinas M and Vaidya AB [9]. Usually, glutamine is the carbon source of TCA cycle but the main source of glutamine is hemoglobin digestion, a process normally reduced in resistant [2]; glutamine can also be used for glutathione production.

On the other hand, several genes implicated in the mitochondrial electron transport chain (ETC) and forming ATP synthase are downregulated [15]. This may indicate that those parasites are worse off producing electron potential across matrix membrane. Interestingly, recent data obtained from a small subset of artemisinin-induced dormant *P. falciparum* parasites show that these parasites need mitochondrial matrix membrane potential to arise from dormancy [35]; the under expression of ETC genes may be associated with dormancy state, previously reported for *P. falciparum* (reviewed in [36])

All of the above suggests that the PfKelch 13 mutant strains of *P. falciparum* would be obliged to constantly synthesize glutathione during this stage to combat the oxidative stress caused by artemisinin. The constant synthesis of glutathione would compel *P. falciparum* to produce sufficient oxoglutarate in the mitochondria from the glucose consumed.

## 6 Conclusions

In sum, we can conclude that the PfCOCP of *P. falciparum* may be playing an important role in response to oxidative stress experienced by the parasite in the presence of artemisinin. PfCOCP allows for the transportation of citrate and oxoglutarate from the matrix to the cytosol for subsequent transformation to glutathione and NADPH productions. This model offers previously unreported metabolic changes in artemisinin-resistant parasites and may represent new therapeutic targets to consider in artemisinin-resistant strains of *P. falciparum*.

## 7 Acknowledgements

To Prof. Ananias Escalante of Temple University (TU) and Prof. Ines Rojas Avendaño of the University of The Andes (ULA) for their accurate comments on the manuscript.

